# Genomic signatures of extreme body size divergence in baboons

**DOI:** 10.1101/578740

**Authors:** Kenneth L. Chiou, Christina M. Bergey, Andrew S. Burrell, Todd R. Disotell, Jeffrey Rogers, Clifford J. Jolly, Jane E. Phillips-Conroy

## Abstract

Kinda and gray-footed chacma baboons occupy opposite extremes of the body size distribution in extant baboons (genus *Papio*). In order to detect signatures of natural selection in these two species, we genotyped 24,790 genome-wide autosomal SNPs from populations of Zambian baboons using double digest RADseq. We scanned the genome for evidence of selection by identifying regions with extreme differentiation between populations. We find evidence of selection on body size influencing multiple genes in one or both species, including *FGF1, ATXN2*, and *PRKCE*. We also find an enriched signal of selection associated with biological processes involved in multicellular organism growth and development, cell proliferation and cell growth, nutrient metabolism, and chondrocyte differentiation. Finally, we find that selection has impacted components of the CCKR signaling pathway, which regulates food intake and metabolism, and the JAK/STAT signaling pathway, which mediates the effect of cytokine signals on processes including epiphyseal chondrocyte proliferation essential for longitudinal bone growth. Our findings highlight promising avenues for future studies disentangling the genetic architecture of body size in primates including humans.

## Introduction

Body size is among the most fundamental variables influencing the diversity of biological traits in living organisms including primates (Haldane 1926; McMahon 1973; Martin 1980; Fleagle 1985). Much of the variation in primate morphology, physiology, ecology, life history, and behavior is intimately linked to variation in body size (Clutton-Brock et al. 1977; Leutenegger & Cheverud 1982; Fleagle 1985). While considerable attention has been dedicated to the proximate hormonal mechanisms underlying body-size variation (Bernstein 2010), few studies (Worley et al. 2014) have examined the genetic basis for body-size variation in nonhuman primates, particularly at the genomic level.

Baboons (genus *Papio*) are well-known and extensively studied cercopithecine primates that exhibit considerable variation in body size, ecology, morphology, and behavior (Jolly 1993; Barrett 2009). The impressive diversity of this genus, combined with a complex evolutionary history following a crown divergence approximately 2.0 – 1.5 million years ago (Zinner et al. 2013; Rogers et al. 2019), have made baboons fruitful models for examining mechanisms of natural selection in wild populations (e.g., Jolly 2001; Henzi & Barrett 2003; Strum 2012).

Kinda (*Papio kindae*) and gray-footed chacma baboons (*P. ursinus griseipes*) are two parapatrically distributed southern African taxa that in some aspects occupy opposing extremes of phenotypic variation in extant baboons. Most strikingly, gray-footed chacma baboons along with other chacma baboons are the largest extant baboons while Kinda baboons are the smallest (Delson et al. 2000; Jolly, Burrell, et al. 2011). The size differences are such that individuals in the normal ranges of variation for the respective species can differ in body weight by a factor of 2.0 or more in males and 1.5 or more in females (Figure 1).The two species differ additionally in a variety of size-related features, including sexual body-size dimorphism and facial length (Jolly, Burrell, et al. 2011). Male Kinda baboons additionally exhibit a gracile, lanky build that lends them an overall juvenile appearance (Jolly, Burrell, et al. 2011). Interestingly, these substantial differences do not represent an insurmountable barrier to successful reproduction, as the two species are involved in ongoing natural hybridization in central Zambia (Jolly, Burrell, et al. 2011).

**Fig. 1:**
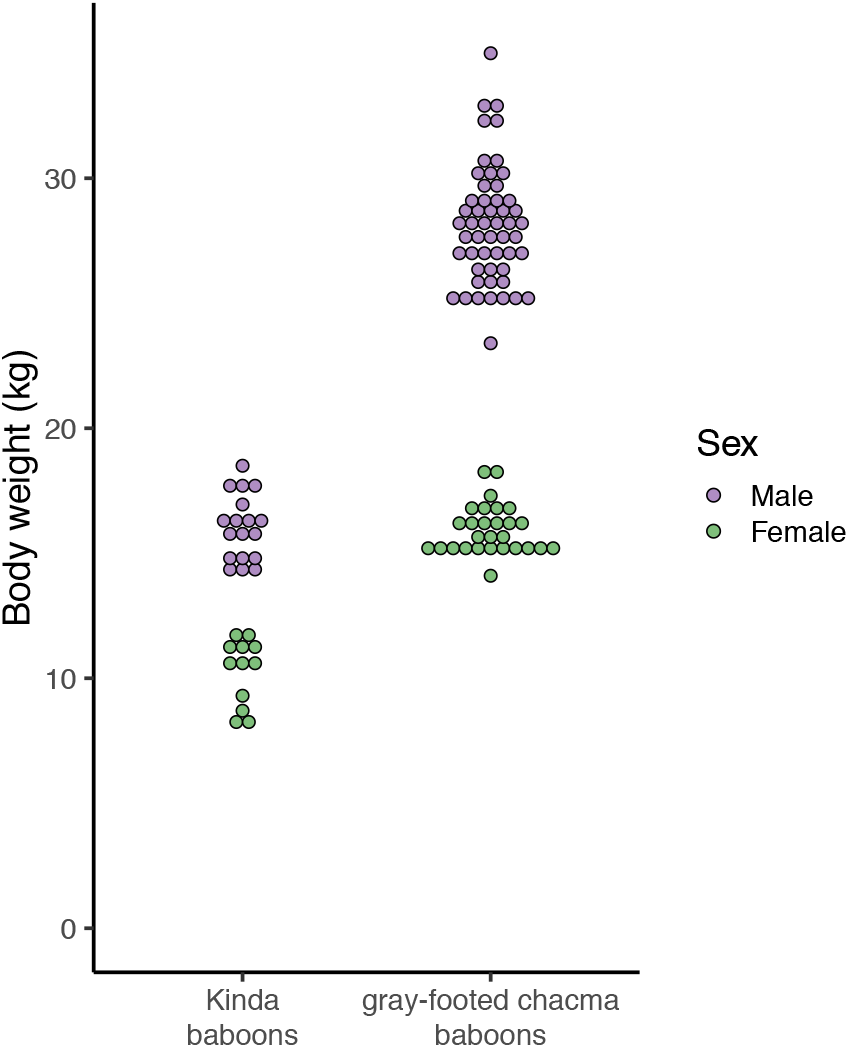
Dot plots comparing body weights of Kinda and gray-footed chacma baboons. All data were obtained from adult individuals live-captured from the wild. Kinda baboons were measured in Kafue National Park, Zambia. Gray-footed chacma baboons were measured in Moremi Game Reserve, Botswana. Body weight measurements from the Moremi baboon population were kindly provided by John Bulger and are used with permission.

In the present study, we genotype Kinda and gray-footed chacma baboons at roughly 25,000 genome-wide polymorphic sites in order to investigate the genetic basis of divergent size-related phenotypes including body mass. We then use a genomic selection-scan approach to identify regions of the genome putatively under selection and use functional and pathway annotations to evaluate our prediction that differentiated genomic loci will exhibit links to growth and metabolism, key traits underlying body size.

## Materials and Methods

### Samples

Genetic material from wild Zambian baboons was derived from blood samples collected in 2011 and 2012 and from fecal samples collected between 2006 and 2007 (Table 1). Sampling was concentrated around Kafue National Park, Zambia, where Kinda and gray-footed chacma baboon distributions join and overlap with evidence of hybridization (Jolly, Burrell, et al. 2011). Blood was drawn from animals that were trapped, tranquilized, then released following established protocols (Brett et al. 1977; Jolly, Phillips-Conroy, et al. 2011). We collected whole blood into evacuated tubes containing EDTA anticoagulant, then fractionated the blood by centrifugation. Plasma, erythrocyte-, and leukocyte-rich fractions were stored immediately in liquid nitrogen and subsequently at −80 °C. A small subset of whole blood was preserved on a Whatman FTA card and stored at room temperature. Feces were obtained noninvasively during field surveys and stored in RNAlater (Ambion) at room temperature, then subsequently at −20 °C. All trapping and sample collection procedures were conducted following laws and regulations in Zambia and with approval from the Animal Studies Committees at Washington University, New York University, and Baylor College of Medicine. This research follows the International Primatological Society’s Code of Best Practices for Field Primatology.

**Table 1:** Samples included in this analysis. Only samples that passed quality and ancestry filters are listed in this table. An expanded list including sample IDs is provided in Table S1.

| Locality | Latitude | Longitude | Year | Ancestry | Tissue type | N |
| --- | --- | --- | --- | --- | --- | --- |
| Chunga | -15.0446° | 25.9994° | 2011 | Kinda | blood | 69 |
| North Nkala Road | -15.9029° | 25.8895° | 2007 | chacma | feces | 1 |
| Ngoma Airstrip | -15.9695° | 25.9376° | 2012 | chacma | blood | 4 |
| Dendro Park | -16.1518° | 26.0599° | 2012 | chacma | blood | 4 |
| Nanzhila Plains | -16.2337° | 25.9612° | 2007 | chacma | feces | 3 |
| Choma | -16.6383° | 27.0308° | 2007 | chacma | feces | 3 |
| Lower Zambezi National Park | -15.9234° | 28.9466° | 2006 | chacma | feces | 5 |

### DNA isolation and library preparation

We sequenced DNA from a total of 129 animals, of which 89 remained after filtering for quality and ancestry as described in the following sections. For blood samples, we extracted DNA from stored leukocyte (*n* = 45), plasma (*n* = 16), or FTA-dried blood spot samples (*n* = 26) using the QIAamp DNA Blood Mini Kit (Qiagen). For fecal samples, we extracted DNA using the QIAamp DNA Stool Mini Kit (Qiagen).

Because fecal DNA is dominated by a high presence of exogenous, mainly microbial DNA, we enriched host DNA from fecal samples using FecalSeq (Chiou & Bergey 2018), which makes use of natural differences in CpG methylation between vertebrate and bacterial genomes to preferentially capture CpG-methylated host DNA from feces. For each sample, we added 150 – 1000 ng of DNA to 1 μl of prepared MBD2-Fc/protein A beads (New England Biolabs), then washed and eluted the DNA as described by Chiou and Bergey (2018). To maximize the fraction of host DNA recovered, we performed two serial rounds of enrichment for each fecal DNA sample.

Due to the high cost of generating whole-genome sequences at the population level, particularly from feces-derived genomes (Snyder-Mackler et al. 2016; Chiou & Bergey 2018), we used double-digest RADseq (ddRAD-seq) (Peterson et al. 2012) to reduce our genomic dataset to a representative and reproducible set of orthologous sites. We prepared multiplexed ddRADseq libraries using *SphI* and *MluCI* and a mean fragment size of 300 bp. If the input DNA quantity was less than 20 ng, as is typically the case with enriched fecal DNA, we used 1 unit of each restriction enzyme. Otherwise, we used 1 – 10 units of each restriction enzyme following a ratio of one unit of enzyme per 20 ng of input DNA. The remainder of the ddRADseq library preparation followed standard ddRADseq procedures with modifications for low quantities when necessary (Chiou & Bergey 2018). We sequenced finished libraries on the Illumina HiSeq 2500 or Illumina HiSeq 4000 platforms using 2 × 100 bp sequencing.

### Sequence alignment and variant identification

Sequencing reads were demultiplexed using sabre (https://github.com/najoshi/sabre), allowing one mismatch, and adapter sequences were trimmed using fastq-mcf (arguments: `-l 15 -q 15 -w 4 -u -P 33`) (Aronesty 2011). We then aligned reads to the anubis baboon reference genome (Panu2.0) (Rogers et al. 2019) using default settings of the Burrows-Wheeler Aligner BWA-MEM algorithm (Li & Durbin 2009; Heng Li 2013). After realignment around indels using the Genome Analysis Toolkit (GATK) (McKenna et al. 2010), we called variants on the complete dataset including all individuals using GATK UnifiedGenotyper (arguments: `–stand_call_conf 50.0 -stand_emit_conf 10.0`) and filtered variant calls using GATK VariantFiltration (filters: `QD < 2.0, MQ < 40.0, FS > 60.0, HaplotypeScore > 13.0, MQRankSum < −12.5, ReadPosRankSum < −8.0`).

Autosomal variants that passed the filters described above were further filtered to exclude variants in repetitive regions and variants with substantial missingness. Variants in repetitive regions were filtered out based on publicly available identifications obtained from UCSC Genome Browser (Meyer et al. 2013) and generated using RepeatMasker (Smit et al. 2015) and Tandem Repeats Finder (Benson 1999) for the baboon reference genome (Panu2.0). After filtering out repetitive regions, 590,666 variants remained with a genotyping rate of 23.1%. We then filtered out variants missing in ≥ 20% of individuals and removed individuals with ≥ 80% missing data using PLINK (`--geno 0.2 --mind 0.8`) (Purcell et al. 2007; Chang et al. 2015). After applying these filters, 24,790 variants and 123 individuals remained, with a genotyping rate of 88.8%.

For ancestry analysis, we created an additionally pruned dataset by removing variants in strong linkage disequilibrium (Bergey et al. 2016). Variants in strong linkage disequilibrium were identified based on sliding windows of 50 SNPs with a step size of 5 SNPs and an *r*^2^ threshold of 0.5 using PLINK (`--indep-pairwise 50 5 0.5`) (Purcell et al. 2007; Chang et al. 2015). The pruned dataset contained 14,642 variants.

### Inference of ancestry

Because the focus of this analysis is on between-species differences and samples were collected from an area containing known hybrids (Jolly, Burrell, et al. 2011), we took the following steps to remove individuals with mixed ancestry from this analysis. We used an unsupervised clustering algorithm implemented in ADMIXTURE to estimate the ancestry of each individual by maximum likelihood (Alexander et al. 2009). For this analysis, we used the pruned dataset of 14,642 variants. We explored through cross validation a range of *K* from 1 to 10 and found that *K* = 2 was the most likely value (Figure S1). We considered individuals to be of pure ancestry if their ancestry estimates inferred by ADMIXTURE were greater than 0.999. By this criterion, 34 individuals of mixed ancestry were excluded from subsequent analysis. Our final dataset contained 69 pure Kinda baboons and 20 pure chacma baboons (Figure S2).

### Estimation of differentiation

Differential selection on populations leads to a significant difference in allele frequencies between populations. The well-known fixation index (*F_ST_*) quantifies this pattern by comparing the variance of allele frequencies within and between populations. Comparatively large values of *F_ST_* at a locus relative to neutral regions indicate a larger differentiation between populations. Regions with extremely high values of *F_ST_* indicate the putative effects of directional selection and are therefore considered to be candidates for selection (Akey et al. 2002).

We first used VCFtools (Danecek et al. 2011) to calculate *F_ST_*, following the equation of Weir and Cockerham (Weir & Cockerham 1984; Cockerham & Weir 1987), for the full dataset of 24,790 SNP loci (including loci in strong linkage disequilibrium). Next, based on the baboon reference genome (Panu2.0) general transfer format (GTF) annotation file obtained from Ensembl (Cunningham et al. 2015), we assigned a single value of *F_ST_* to each gene calculated as the average *F_ST_* of all SNPs falling within the boundaries of a protein-coding gene ±50 bp.

We used a permutation test to assess the significance of *F_ST_* estimates for each gene. In each iteration, we randomly reassigned *F_ST_* values to SNPs by sampling with replacement from all *F_ST_* values in the SNP dataset. *F_ST_* values were then assigned to genes following the procedure described above. Sampling was conducted first for 10,000 replicates. A *p_Fst_* value was then calculated as the proportion of *F_ST_* values in the randomized samples that were greater than the actual *F_ST_* value for each gene. If significance (*p* < 0.05) could not be confidently determined after 10,000 replicates, we allowed the analysis for remaining genes to continue to a maximum of 100 million replicates, after which all *p_Fst_* values could be confidently distinguished from a significance threshold of α = 0.05. This procedure increases computational efficiency by focusing computational effort on genes whose significance is more difficult to determine.

### Functional enrichment analysis

We next conducted enrichment analyses to test for overall patterns of differentiation in genes classified according to known functions or roles in biological pathways.

To test for enrichment of Gene Ontology (GO) (Gene Ontology Consortium 2000, 2015) terms, we first downloaded GO annotations for all proteins in the baboon reference genome (Panu2.0) using biomaRt (Durinck et al. 2009). We limited terms to those in the “biological process” ontology with a minimum of 10 annotated genes in our dataset. In order to test for GO terms associated with significantly shifts in the distribution of *p_Fst_*, we ran a modified Kolmogorov–Smirnov test implemented in the R package topGO (Alexa & Rahnenführer 2018). We ran the test using the “weight01” algorithm, which increases biological signal by decorrelating the underlying GO graph structure (Alexa et al. 2006).

To test for enrichment of PANTHER pathways (Mi, Muruganujan & Thomas 2013; Mi, Muruganujan, Casagrande, et al. 2013), we first downloaded PANTHER annotations for the rhesus macaque genome due to the current unavailability of baboon gene sets in PANTHER. We next assigned baboon proteins to rhesus macaque proteins by homology using HMMER3 phmmer (Finn et al. 2011). We excluded matches with E-values > 0.0001 and retained a single protein with the lowest E-value. We then assigned PANTHER pathways based on UniProt identifications in the complete PANTHER sequence classifications for the rhesus macaque proteome. We excluded duplicate assignment of PANTHER pathways to genes in order to avoid redundancy. In order to test for PANTHER pathways associated with extremely differentiated genes, we conducted one-sided Wilcoxon rank sum tests following Mi et al. (2013), evaluating the alternative hypothesis that *p_Fst_* values of genes assigned to a pathway were lower than *p_Fst_* values not assigned to that pathway. To test for robustness of the analysis to filtering criteria, enrichment tests were run using threshold values (*n*) ranging from 2 to 5, each time filtering the dataset to include only pathways represented by at least *n* genes. All enrichment analyses were run in R (R Core Team 2013).

For both GO and PANTHER enrichment analyses, terms and pathways with a *p*-value < 0.05 were considered enriched. Correction for multiple hypothesis testing was not conducted due to a lack of statistical power and the exploratory nature of these analyses.

## Results

From the 89 Zambian baboons in our ancestry-filtered dataset (69 Kinda and 20 gray-footed chacma baboons), we genotyped 24,790 SNPs and used a genome-wide selection scan approach based on *F_ST_* to identify candidate genes showing signatures of high differentiation between populations (Figure S3). Using annotation information from the baboon reference genome, we retained genes which included genotyped SNPs falling within or near their boundaries. Out of 27,857 genes in the baboon reference genome, we thus derived a sample of 2,213 genes, or approximately 8% of all genes. The large fraction of genes unrepresented in our genic dataset reflects a combination of 1) the purposeful shared enrichment of a small fraction of randomly sampled regions of the genome characteristic of double-digest RADseq (Peterson et al. 2012) as well as 2) the relative low-coverage of libraries derived from feces. For each gene in this dataset, we next assigned a mean *F_ST_* and identified through permutation 175 candidate genes with extremely high *F_ST_*(*p_Fst_* < 0.05; Table S2). After controlling for multiple hypothesis testing using the false discovery rate (FDR) (Benjamini & Hochberg 1995), we identified 10 genes considered to be especially strong candidates for differentiation (FDR < 0.05) (Table 2).

**Table 2:**
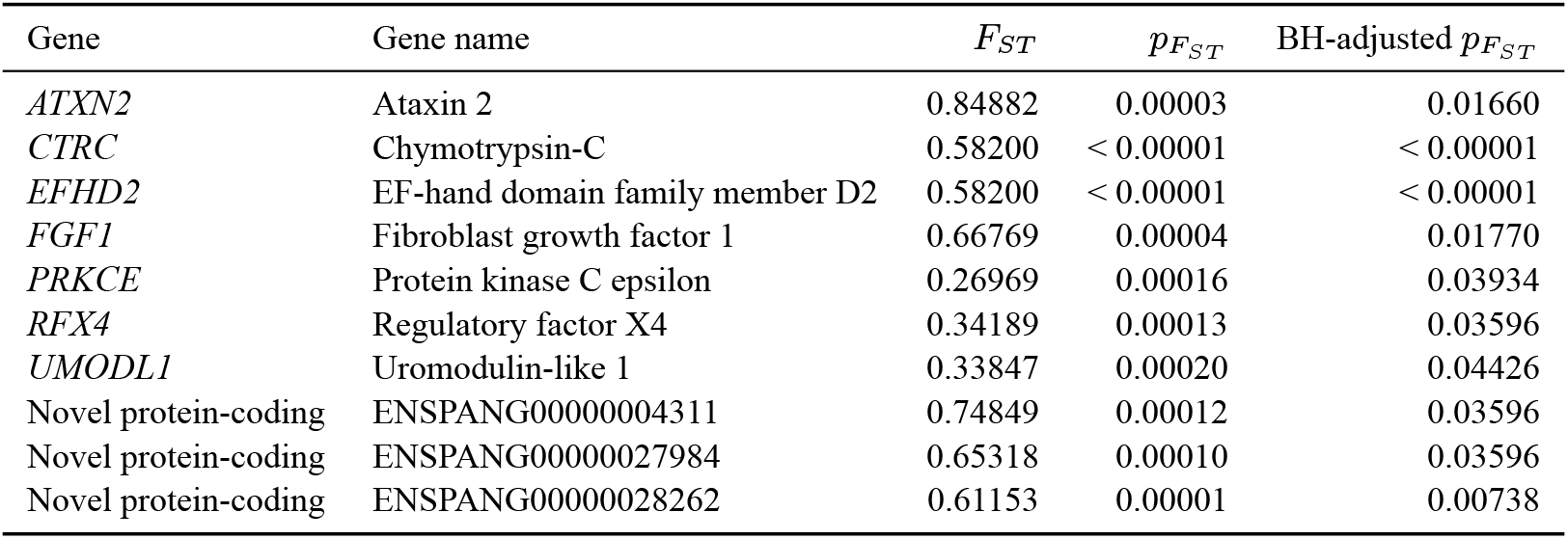
Genes that are significantly differentiated between Kinda and chacma baboon populations based on *F_ST_*. Genes listed here were significant after controlling for multiple testing (FDR < 0.05) and are therefore considered to be strong candidates for differentiation.

Three of the 10 genes are novel protein-coding genes lacking information about function. We assessed the remaining genes for functions that are potentially relevant in producing the morphologically and behaviorally divergent Kinda and chacma baboon phenotypes.

Several genes have known functions related to developmental or metabolic processes. One of them, fibroblast growth factor 1 (*FGF1*), promotes cell division in a variety of mesoderm- and neuroectoderm-derived cells, and also promotes angiogenesis (Chiu et al. 1990; Nabel et al. 1993) and metabolic homeostasis (Jonker et al. 2012). It is also expressed in proliferating and hypertrophic chondrocytes in human fetal growth plates, with inhibitory effects *in vitro* on chondrocyte proliferation (Krejci et al. 2007). Another outlier gene, ataxin 2 (*ATXN2*) has been associated with weight gain and insulin resistance in mice (Kiehl et al. 2006; Lastres-Becker et al. 2008) and hypertension and diabetes mellitus in humans (Levy et al. 2009; Lastres-Becker et al. 2016). Evidence from both yeast and mouse models indicates that ATXN2 blunts mechanistic target of rapamycin (mTOR) signaling (Fittschen et al. 2015; Lastres-Becker et al. 2016; Meierhofer et al. 2016), suggesting that it functions as a modulator of nutrition, cell growth, and metabolism. Finally, protein kinase C epsilon (*PRKCE*) and its encoded protein PKCε is a target of mTOR signaling through the protein complex mTORC2 and has been reported to play a role in diverse signaling mechanisms including cell proliferation, differentiation, gene expression, and metabolism (Akita 2002). It also plays a role in regulating the secretion of insulin in response to glucose signals in pancreatic β cells (Warwar et al. 2008). Interestingly, a *de novo* constitutional mutation in *PRKCE* linked to impaired mTORC2 signaling was found in a clinical case of SHORT syndrome, a condition characterized by short stature, intrauterine growth restriction, and insulin resistance (Alcantara et al. 2017).

Two genes are relatively little known but appear to play a role in sex-specific developmental processes. Uromodulin-like 1 (*UMODL1*) helps regulate ovarian development in mice (Wang et al. 2012). Regulatory factor X4 (*RFX4*) is highly expressed in the central nervous system in mice and may mediate ciliogenesis by modulating Sonic hedgehog (Shh) signaling during development (Ashique et al. 2009).

Interestingly, *RFX4* in humans is expressed entirely in the testis (Morotomi-Yano et al. 2002). Testicular morphology and physiology have a well-known association with taxon-specific social behavior and levels of sperm competition in mammals (Harcourt et al. 1981; Møller 1988, 1989; Dixson & Anderson 2004) and baboons in particular (Jolly & Phillips-Conroy 2003, 2006). Given considerable differences between Kinda and chacma baboons in both size and behavior, differentiation of *RFX4* suggests a possible proximate mechanism. Closer inspection of SNPs in *RFX4* indicates that, in the most highly differentiated SNP located on chromosome 11 at position 105851669 on the baboon reference genome (Panu2.0, derived from an anubis baboon, *Papio anubis*) (Rogers et al. 2019), the high-frequency allele in chacma baboons is shared with both the anubis baboon reference individual and the rhesus macaque reference individual as an outgroup (Table S3), indicating that the high distinctiveness of *RFX4* may be driven by directional selection on Kinda baboons. These interpretations remain speculative, however, in the absence of additional sequence and expression information from *RFX4*, as well as functional validation.

Two genes have been linked to pathways related to behavior. EF-hand domain family member D2 (*EFHD2*) plays a role in immune and brain cell function (Dütting et al. 2011). While it is best-known for its role in tau-related neurodegenerative diseases including Alzheimer’s disease (Vega 2016), it has also been implicated in pathways associated with aggression-related behavior (Malki et al. 2014). *PRKCE* knockout mice show reduced anxiety-like behaviors and reduced levels of the stress hormones corticosterone and adrenocorticotrophic hormone (Hodge et al. 2002).

In order to analyze the functional processes associated with highly differentiated genes, we conducted a Gene Ontology enrichment analysis using methods that take into account the correlated underlying graph topology of GO terms (Alexa et al. 2006; Alexa & Rahnenführer 2018). These analyses identified 66 enriched GO terms (Table S4), indicating that they were associated with higher genome-wide differentiation than expected by chance (*p* < 0.05). Correction for multiple hypothesis testing was not applied due to limited statistical power as well as the exploratory nature of the analyses. Because of the heightened risk of false positives, we treat individual results with skepticism but report on overall patterns in the data.

The GO analyses revealed the enrichment of several terms broadly related to organismal growth, including “multicellular organism growth” (*p* = 0.015), “developmental growth” (*p* = 0.008), and “positive regulation of developmental growth” (*p* = 0.001). Other significantly enriched terms were related to the mitotic process (e.g., “negative regulation of mitotic nuclear division” [*p* = 0.001], “negative regulation of mitotic cell cycle phase transition” [*p* = 0.030], and “mitotic sister chromatid segregation” [*p* = 0.031]), cell proliferation (e.g., “regulation of endothelial cell proliferation” [*p* = 0.004] and “negative regulation of epithelial cell proliferation” [*p* = 0.004]), or cell growth (e.g., “regulation of cell size” [*p* = 0.032] and “positive regulation of cell growth” [*p* = 0.035]). Importantly, changes to both cell division (affecting cell number) and cell growth (affecting cell size) underlie body size variation in multicellular organisms (Schmelzle & Hall 2000).

Our analyses revealed the enrichment of several terms related to nutrient metabolism and response to environmental inputs. These terms included “response to regulation of glucose metabolic process” (*p* = 0.038), “regulation of insulin secretion” (*p* = 0.047), and “response to nutrient levels” (*p* = 0.004). We also found the enrichment of one term, “chondrocyte differentiation” (*p* = 0.009), reflecting the critical role played by chondrocytes in developmental bone growth. Other enriched terms potentially related to differences in body size included “connective tissue development” (*p* = 0.002) and “regulation of fat cell differentiation” (*p* < 0.001).

We next assessed pathway annotations using the PANTHER classification system (Mi, Muruganujan & Thomas 2013; Mi, Muruganujan, Casagrande, et al. 2013) to identify pathways with enriched differentiation between Kinda and chacma baboons. These analyses identified only two enriched PANTHER pathways (Table 3), indicating that they were associated with higher genome-wide differentiation than expected by chance (*p* < 0.05). These pathways were the CCKR signaling pathway and the JAK/STAT signaling pathway. As with the GO enrichment analysis, correction for multiple testing was not conducted for this exploratory analysis.

**Table 3:**
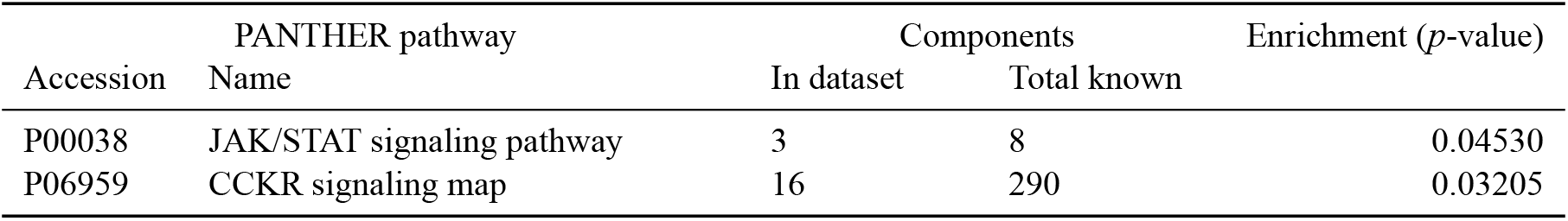
Pathways with enriched differentiation. Pathways that are enriched exhibit an overall shift in *p_F_ST__*. We considered enrichment of pathways to be significant if we found *p* < 0.05.

| Accession | PANTHER pathway<br>Name | Components | | Enrichment ($p$-value) |
| --- | --- | --- | --- | --- |
|  |  | In dataset | Total known |  |
| P00038 | JAK/STAT signaling pathway | 3 | 8 | 0.04530 |
| P06959 | CCKR signaling map | 16 | 290 | 0.03205 |

Out of fourteen genes in our dataset assigned to the CCKR signaling pathway, only *PRKCE* was significantly differentiated after correcting for multiple hypothesis testing (Table S5). An additional three genes passed a threshold of *p_Fst_* < 0.05 prior to correction for multiple testing.

Out of four genes in our dataset assigned to the JAK/STAT pathway, none was significantly differentiated despite enrichment of the overall pathway (Table S6). One gene (*STAT3*) was shared with the CCKR signaling pathway. The most differentiated SNP was found in *JAK1* on chromosome 1 at position 66,951,082 on the baboon reference genome, which is located downstream of the protein-coding sequence. Interestingly, at this SNP, the alternate allele occurs at high frequency in Kinda baboons and is shared with a rhesus macaque outgroup, while the anubis baboon reference allele is found at high frequency in chacma baboons (Table S6). *JAK1*-knockout mice are associated with significantly smaller perinatal body masses (Rodig et al. 1998; Liu et al. 2004). These results suggest a possible role for JAK/STAT signaling in divergent body size phenotypes of Kinda and chacma baboons. The mechanisms and the genes involved, however, remain unknown.

## Discussion

Using genome-wide polymorphism data from Zambian baboons occupying opposite extremes of the extant baboon size distribution, we were able to identify putatively selected genes that may underlie divergent phenotypes in the two species, particularly those related to body size. We identified putative selection at three levels: genes, biological processes, and biological pathways.

Of the ten genes classified as possible targets of selection based on significant differentiation after controlling the false discovery rate (FDR < 0.05), several are known to be involved in growth or metabolic processes and may contribute to differences in body size. Intriguingly, a number of these genes are also linked to targets or regulators of the interconnected insulin/insulin-like growth factor (IIS) and mechanistic target of rapamycin (mTOR) pathways, which constitute two of the most important regulators of organism growth in response to nutrient availability in metazoans (Oldham & Hafen 2003; Saxton & Sabatini 2017; Koyama & Mirth 2018). *FGF1*, for instance, has been shown to play a critical role in nutrient homeostasis, with FGF1-knockout mice developing insulin resistance when stressed with a high-fat diet (Jonker et al. 2012). *ATXN2* has been shown to modulate nutrition and metabolism by blunting mTOR signaling, with ATXN2-knockout mice exhibiting increased mTOR-dependent ribosomal S6 phosphorylation (Fittschen et al. 2015). PKCε is targeted by mTORC2 and appears to play a critical role in activation of Akt (Alcantara et al. 2017), a key effector of insulin/PI3K signaling (Saxton & Sabatini 2017).

Through functional enrichment analyses using GO terms from the biological processes ontology, we found robust evidence that terms related to body size differences were enriched with high *F_ST_*, indicating the associated gene sets as a whole were associated with high differentiation even if few or no individual genes were individually identified as candidate genes. Enriched categories included high-level terms such as “multicellular organism growth” and “developmental growth”, as well as terms related to control of cell division and cell growth. We also found the enrichment of several terms related to nutrient signaling and nutrient metabolism, as well as of one term related to chondrocyte differentiation.

Through a similar analysis of biological pathways using PANTHER classifications, we found that only two pathways, the CCKR signaling pathway and the JAK/STAT signaling pathway, were enriched with high *F_ST_*.

The CCKR signaling pathway coordinates food intake by controlling visceral functions and behavior (Ritter et al. 1999; Woods 2005). Cholecystokinin (CCK) stimulates the secretion of pancreatic digestive enzymes, the production and excretion of bile into the intestine, the delay of gastric emptying, and the inhibition of food intake (Ritter et al. 1999; Rehfeld 2017). Through its role as a satiety signal and regulator of energy intake, CCK is thought to act as a regulator of body weight (Ritter et al. 1999; Woods 2005). It also appears to act synergistically with leptin to regulate body weight even absent differences in food intake (Matson & Ritter 1999; Matson et al. 2000). Evidence for CCK underlying the evolution of body size across populations or species, however, remains sparse.

The JAK/STAT signaling pathway is a major cell-signaling mechanism, originating prior to the divergence of protostomes and deuterostomes (Liongue & Ward 2013). Cellular responses to myriad hormones, growth factors, and cytokines are mediated by this evolutionarily conserved pathway (Kisseleva et al. 2002). JAK/-STAT signaling is capable of eliciting specialized responses to extracellular signals that depend on the signal molecule and the tissue or cellular context. Predictably, mutations that affect JAK/STAT signaling, for instance by suppressing its components or by constitutively activating or failing to regulate its activity, also affect these processes and thereby influence organismal phenotypes (O’Shea et al. 2002; Rawlings et al. 2004).

While JAK/STAT signaling underlies diverse responses including mammary gland development, hematopoiesis, and immune cell development (Watson & Burdon 1996; Ward et al. 2000; Ghoreschi et al. 2009), the statistical enrichment of this PANTHER pathway is particularly intriguing due to its known function in organism growth (Yakar et al. 2018) and metabolism (Dodington et al. 2018). Knockout experiments in mouse models demonstrate the impacts of multiple JAKs and STATs on bone development and body mass (Jiliang Li 2013). JAK1-deficient mice, for instance, exhibit significantly smaller perinatal body mass than their littermates (Rodig et al. 1998). The selective inactivation of STAT3 in osteoblasts is also associated with lower body mass and bone mass (Zhou et al. 2011). JAK/STAT signaling additionally has profound impacts on adipose tissue development and physiology (Richard & Stephens 2014), although as previously mentioned, the relevance of these functions is at present unclear due to a paucity of morphometric data on adiposity.

Notably, JAK/STAT is known to serve as a signaling mechanism for the GH/IGF1 axis, which is one of the major endocrine systems regulating somatic growth and stature (Yakar et al. 2018). Growth hormone (GH) secreted in the anterior pituitary stimulates the production of insulin-like growth factor 1 (IGF1) through JAK/STAT signals. Together, GH and IGF1 play essential roles in regulating bone growth through chondrocyte proliferation at epiphyseal growth plates (Yakar et al. 2018). These functions can be impacted through modifications to JAK/STAT signaling. Suppressor of cytokine signaling 2 (SOCS2), for instance, is a negative regulator of GH action through inhibition of JAK/STAT, and SOCS2 deficiency in mice is associated with gigantism and increased longitudinal bone growth (Metcalf et al. 2000; Pass et al. 2012).

JAK/STAT function on chondrocyte proliferation has also been shown to interact with fibroblast growth factor (FGF) signaling. FGF1, which was identified in this study as a strong candidate for differentiation as discussed above, is expressed in human fetal growth plate cartilage (Krejci et al. 2007) and has been shown to suppress chondrocyte proliferation in mouse and rat cells through induction of STAT1, which has anti-proliferative functions (Sahni et al. 1999).

While our combined analyses suggest several candidate genes and gene sets involved in extreme body size divergence in baboons, our analyses are by no means comprehensive due to methodological and statistical constraints. Our use of double-digest RADseq, for instance, increased our read coverage and decreased the cost of sequencing relative to whole-genome sequencing (Peterson et al. 2012) but at the expense of limiting the fraction of the genome available for this analysis (Lowry et al. 2017). We therefore expect to have missed many loci under selection, as well as experiencing decreased overall statistical power for detecting differentiated loci in our dataset. Furthermore, as a complex trait, body size is controlled by a large number of genes, most of which have such small effect sizes that they are unlikely to be detected by genomic scans (Yang et al. 2010; Boyle et al. 2017).

Despite these limitations, our analyses highlight genetic differences between two baboon species that differ greatly in body size, which may therefore lend insights into both directional selection and the genetic architecture of organismal size. Intriguingly, we found a signal of body size divergence at multiple biological levels: genes, biological processes, and biological pathways. Given the small proportion of the genome available for analysis, the consistent signal of body size selection across analyses reinforces our conclusion that profound body size differences between these two species have a broad genetic basis.

Additional data are needed to incorporate a higher degree of sequence information and to assess functional impacts of coding as well as regulatory SNPs. These data will enable the generation of more detailed hypotheses that can subsequently be evaluated using functional experimentation. Assessment of additional gene sequences, for instance using whole-genome shotgun sequencing or exome capture methodologies, will incorporate genes that were not represented in our reduced-representation genome dataset. Apart from the highly likely possibility of finding additional candidate genes, these data will also enable more detailed and higher-power analyses, which would for instance allow for corrections for multiple testing that were not practical for this study.

Despite the extreme size difference between Kinda and chacma baboons, these species hybridize in Zambia (Jolly, Burrell, et al. 2011), providing an additional opportunity to disentangle the genetic architecture of body size. Genome-wide association studies of hybrids offers a particularly promising approach. Analysis of hybrids may also lend important insights into selection on variants underlying differences in body size through their degree and directionality of introgression.

## Acknowledgments

We thank the Zambia Wildlife Authority (now the Department of National Parks & Wildlife) and the University of Zambia for granting permission and providing support for fieldwork. We thank John Bulger for sharing body weight data from the Moremi baboon population. This work was supported by the National Science Foundation [grant numbers BCS 1341018, SMA 1338524, BCS 1029302, BCS 1029323, and BCS 1029451]; the Leakey Foundation; and the National Geographic Society. The Genome Technology Center at New York University is supported by the National Center for Advancing Translational Sciences at the National Institutes of Health [UL1 TR00038] and the National Cancer Institute at the National Institutes of Health [P30 CA016087]. K.L.C. is supported by the National Institute of Aging at the National Institutes of Health [T32 AG000057].

## Data Accessibility

Sequence reads are deposited in the NCBI Sequence Read Archive (SRA) under BioProject PRJNA486659, with SRA accession numbers SRR7717274-SRR7717402. All code for this project is available on GitHub at https://github.com/kchiou/kafue-baboons-ddrad.

## Author Contributions

K.L.C., C.M.B., A.S.B., T.R.D., J.R., C.J.J., and J.P.C. designed the research and collected samples. K.L.C. performed the research. K.L.C. and C.M.B. analyzed the data. K.L.C. wrote the paper. All authors read, revised, and approved the manuscript.

