## Supplemental Figures and Tables for "Genomic signatures of extreme body size divergence in baboons"

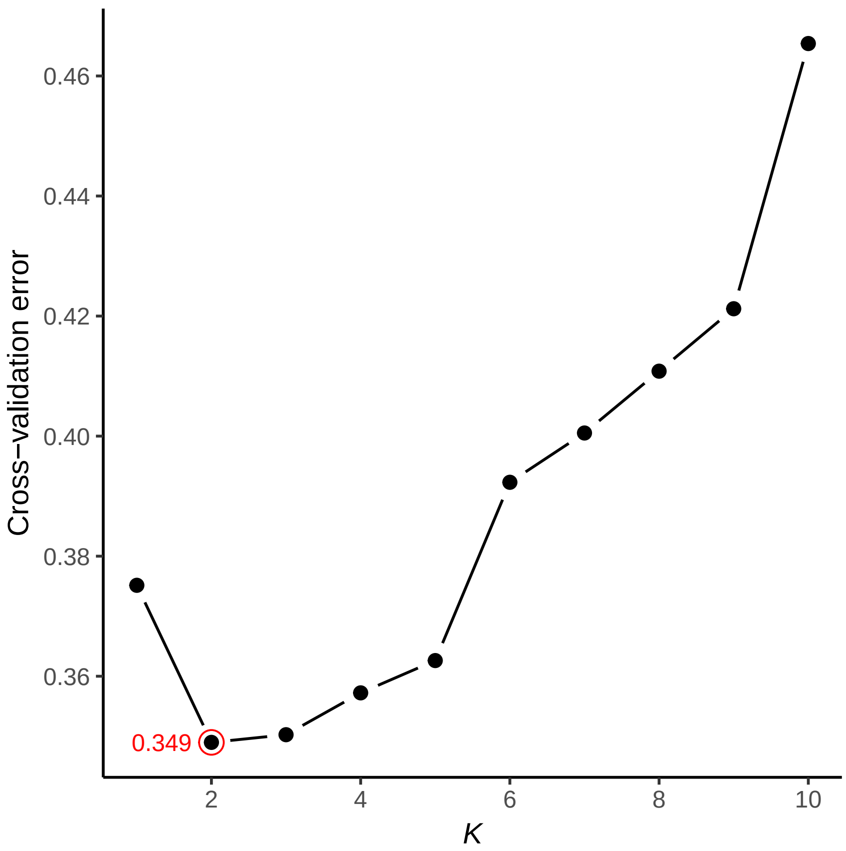


Supplementary Fig. S1: Cross-validated error results from ADMIXTURE runs in which $K$ varied from 1 to 10.

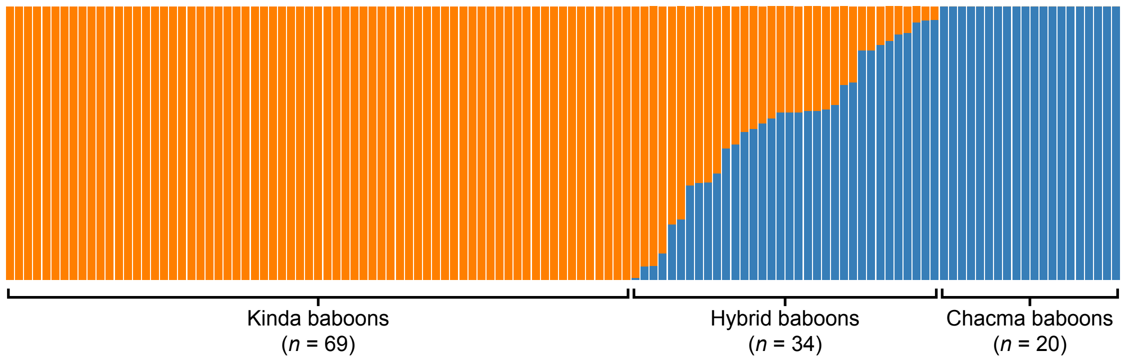


Supplementary Fig. S2: Taxonomic assignment based on ancestry estimates calculated using ADMIXTURE. Of the 129 animals sequenced, 6 were filtered out of the analysis due to excessive missingness and are not shown here.

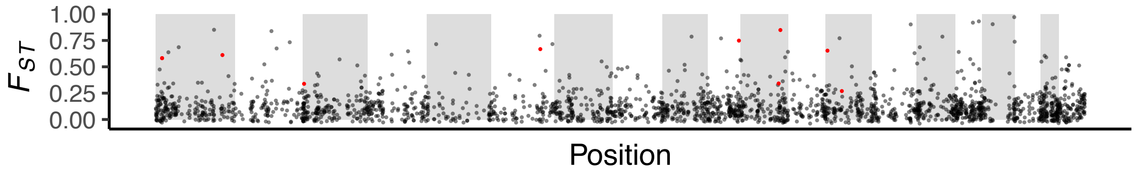


Supplementary Fig. S3: Distribution of $F_{ST}$ for protein-coding genes. Genes identified as significant by permutation are displayed in red (FDR < 0.05). Background shading indicates the position of the autosomal chromosomes in order (i.e., chromosomes 1 - 20).

Supplementary Table S1: Full list of samples included in this analysis. Samples that were sequenced but failed quality-control filters or exhibited hybrid ancestry are not included in this table.

| Sample ID | Tissue type | Locality | Ancestry | SRA Accession |
|---|---|---|---|---|
| BZ11-001 | leukocyte | Chunga | Kinda | SRR7717396 |
| BZ11-002 | leukocyte | Chunga | Kinda | SRR7717393 |
| BZ11-003 | leukocyte | Chunga | Kinda | SRR7717394 |
| BZ11-004 | leukocyte | Chunga | Kinda | SRR7717399 |
| BZ11-005 | leukocyte | Chunga | Kinda | SRR7717400 |
| BZ11-006 | leukocyte | Chunga | Kinda | SRR7717397 |
| BZ11-007 | leukocyte | Chunga | Kinda | SRR7717398 |
| BZ11-008 | leukocyte | Chunga | Kinda | SRR7717401 |
| BZ11-009 | leukocyte | Chunga | Kinda | SRR7717402 |
| BZ11-010 | leukocyte | Chunga | Kinda | SRR7717384 |
| BZ11-011 | leukocyte | Chunga | Kinda | SRR7717383 |
| BZ11-012 | leukocyte | Chunga | Kinda | SRR7717386 |
| BZ11-013 | FTA blood spot | Chunga | Kinda | SRR7717385 |
| BZ11-014 | leukocyte | Chunga | Kinda | SRR7717388 |
| BZ11-015 | leukocyte | Chunga | Kinda | SRR7717387 |
| BZ11-016 | leukocyte | Chunga | Kinda | SRR7717390 |
| BZ11-017 | leukocyte | Chunga | Kinda | SRR7717389 |
| BZ11-018 | leukocyte | Chunga | Kinda | SRR7717392 |
| BZ11-019 | leukocyte | Chunga | Kinda | SRR7717391 |
| BZ11-020 | leukocyte | Chunga | Kinda | SRR7717293 |
| BZ11-021 | FTA blood spot | Chunga | Kinda | SRR7717294 |
| BZ11-022 | FTA blood spot | Chunga | Kinda | SRR7717295 |
| BZ11-023 | FTA blood spot | Chunga | Kinda | SRR7717296 |
| BZ11-024 | leukocyte | Chunga | Kinda | SRR7717297 |
| BZ11-025 | leukocyte | Chunga | Kinda | SRR7717298 |
| BZ11-026 | FTA blood spot | Chunga | Kinda | SRR7717299 |
| BZ11-028 | leukocyte | Chunga | Kinda | SRR7717300 |
| BZ11-029 | leukocyte | Chunga | Kinda | SRR7717301 |
| BZ11-030 | leukocyte | Chunga | Kinda | SRR7717302 |
| BZ11-031 | leukocyte | Chunga | Kinda | SRR7717281 |
| BZ11-032 | leukocyte | Chunga | Kinda | SRR7717280 |
| BZ11-033 | leukocyte | Chunga | Kinda | SRR7717279 |
| BZ11-034 | leukocyte | Chunga | Kinda | SRR7717278 |
| BZ11-035 | leukocyte | Chunga | Kinda | SRR7717277 |
| BZ11-036 | leukocyte | Chunga | Kinda | SRR7717276 |
| BZ11-037 | leukocyte | Chunga | Kinda | SRR7717275 |
| BZ11-038 | leukocyte | Chunga | Kinda | SRR7717274 |
| BZ11-039 | leukocyte | Chunga | Kinda | SRR7717283 |
| BZ11-040 | leukocyte | Chunga | Kinda | SRR7717282 |
| BZ11-041 | leukocyte | Chunga | Kinda | SRR7717321 |
| BZ11-042 | leukocyte | Chunga | Kinda | SRR7717322 |
| BZ11-043 | FTA blood spot | Chunga | Kinda | SRR7717319 |
| BZ11-045 | leukocyte | Chunga | Kinda | SRR7717320 |
| BZ11-046 | FTA blood spot | Chunga | Kinda | SRR7717317 |
| BZ11-047 | leukocyte | Chunga | Kinda | SRR7717318 |
| BZ11-048 | FTA blood spot | Chunga | Kinda | SRR7717315 |
| BZ11-050 | leukocyte | Chunga | Kinda | SRR7717316 |
| BZ11-051 | FTA blood spot | Chunga | Kinda | SRR7717313 |
| BZ11-052 | FTA blood spot | Chunga | Kinda | SRR7717314 |
| BZ11-053 | leukocyte | Chunga | Kinda | SRR7717310 |
| BZ11-054 | leukocyte | Chunga | Kinda | SRR7717309 |
| BZ11-056 | leukocyte | Chunga | Kinda | SRR7717312 |

continued on next page

Supplementary Table S1 – continued from previous page

| Sample ID | Tissue type | Locality | Ancestry | SRA Accession |
|---|---|---|---|---|
| BZ11-058 | leukocyte | Chunga | Kinda | SRR7717306 |
| BZ11-059 | leukocyte | Chunga | Kinda | SRR7717305 |
| BZ11-061 | FTA blood spot | Chunga | Kinda | SRR7717308 |
| BZ11-062 | FTA blood spot | Chunga | Kinda | SRR7717307 |
| BZ11-063 | FTA blood spot | Chunga | Kinda | SRR7717304 |
| BZ11-064 | FTA blood spot | Chunga | Kinda | SRR7717303 |
| BZ11-065 | FTA blood spot | Chunga | Kinda | SRR7717368 |
| BZ11-066 | FTA blood spot | Chunga | Kinda | SRR7717369 |
| BZ11-067 | FTA blood spot | Chunga | Kinda | SRR7717370 |
| BZ11-068 | FTA blood spot | Chunga | Kinda | SRR7717371 |
| BZ11-070 | FTA blood spot | Chunga | Kinda | SRR7717365 |
| BZ11-071 | FTA blood spot | Chunga | Kinda | SRR7717366 |
| BZ11-072 | FTA blood spot | Chunga | Kinda | SRR7717367 |
| BZ11-073 | FTA blood spot | Chunga | Kinda | SRR7717361 |
| BZ11-074 | FTA blood spot | Chunga | Kinda | SRR7717362 |
| BZ11-075 | FTA blood spot | Chunga | Kinda | SRR7717340 |
| BZ11-076 | FTA blood spot | Chunga | Kinda | SRR7717334 |
| BZ07-042 | feces | North Nkala Road | chacma | SRR7717329 |
| BZ12-003 | plasma | Ngoma Airstrip | chacma | SRR7717341 |
| BZ12-006 | plasma | Ngoma Airstrip | chacma | SRR7717338 |
| BZ12-008 | plasma | Ngoma Airstrip | chacma | SRR7717360 |
| BZ12-009 | plasma | Ngoma Airstrip | chacma | SRR7717344 |
| BZ12-030 | plasma | Dendro Park | chacma | SRR7717348 |
| BZ12-031 | plasma | Dendro Park | chacma | SRR7717349 |
| BZ12-032 | plasma | Dendro Park | chacma | SRR7717346 |
| BZ12-033 | plasma | Dendro Park | chacma | SRR7717347 |
| BZ07-029 | feces | Nanzhila Plains | chacma | SRR7717328 |
| BZ07-032 | feces | Nanzhila Plains | chacma | SRR7717326 |
| BZ07-034 | feces | Nanzhila Plains | chacma | SRR7717325 |
| BZ07-004 | feces | Choma | chacma | SRR7717356 |
| BZ07-005 | feces | Choma | chacma | SRR7717359 |
| BZ07-007 | feces | Choma | chacma | SRR7717358 |
| BZ06-218 | feces | Lower Zambezi National Park | chacma | SRR7717351 |
| BZ06-220 | feces | Lower Zambezi National Park | chacma | SRR7717350 |
| BZ06-221 | feces | Lower Zambezi National Park | chacma | SRR7717353 |
| BZ06-225 | feces | Lower Zambezi National Park | chacma | SRR7717355 |
| BZ06-227 | feces | Lower Zambezi National Park | chacma | SRR7717354 |

Supplementary Table S2: Full list of genes with significant $F_{ST}$ ($p_{F_{ST}}$ < 0.05) prior to correction for multiple testing.

| Gene | $F_{ST}$ | $p_{F_{ST}}$ |
|---|---|---|
| *ABCC6* | 0.29510 | 0.03379 |
| *ADAM19* | 0.46159 | 0.04911 |
| *AFDN* | 0.31298 | 0.01436 |
| *AHCTF1* | 0.41161 | 0.02608 |
| *ALDH7A1* | 0.32752 | 0.03378 |
| *ALLC* | 0.31691 | 0.02241 |
| *ALMS1* | 0.40316 | 0.00149 |
| *ALPK2* | 0.23213 | 0.03372 |
| *AQP7* | 0.31983 | 0.02067 |
| *ARCN1* | 0.26019 | 0.00734 |
| *ATP9B* | 0.26107 | 0.02414 |
| *ATXN2* | 0.84882 | 0.00003 |
| *BCAS3* | 0.57289 | 0.02749 |
| *BCL9L* | 0.90226 | 0.00153 |
| *BMP7* | 0.25936 | 0.00059 |
| *C16orf62* | 0.36272 | 0.00939 |
| *CACNA1D* | 0.52348 | 0.03682 |
| *CACNA2D4* | 0.55962 | 0.02934 |
| *CAPN9* | 0.27515 | 0.01603 |
| *CD226* | 0.24492 | 0.00979 |
| *CFAP46* | 0.39592 | 0.03196 |
| *CHST11* | 0.32745 | 0.00345 |
| *CIB3* | 0.30536 | 0.02744 |
| *COL27A1* | 0.57369 | 0.00249 |
| *COPG2* | 0.51315 | 0.03892 |
| *CTRC* | 0.58200 | < 0.00001 |
| *DENND6B* | 0.56703 | 0.00256 |
| *DIS3L2* | 0.27283 | 0.02388 |
| *DNA2* | 0.46812 | 0.04726 |
| *DNER* | 0.27585 | 0.04829 |
| *DPP6* | 0.20804 | 0.04012 |
| *ECE2* | 0.83854 | 0.00330 |
| *EDIL3* | 0.37468 | 0.04026 |
| *EFHD2* | 0.58200 | < 0.00001 |
| *EHD2* | 0.39810 | 0.03011 |
| *EP300* | 0.41123 | 0.02664 |
| *EPB41L4B* | 0.38656 | 0.03501 |
| *ESRRB* | 0.24742 | 0.04475 |
| *ETV7* | 0.39259 | 0.03271 |
| *FAF2* | 0.71554 | 0.01160 |
| *FAM149A* | 0.24276 | 0.00791 |
| *FAM210A* | 0.73778 | 0.01023 |
| *FAM3D* | 0.31282 | 0.02341 |
| *FBRSL1* | 0.64186 | 0.01737 |
| *FBXO10* | 0.28294 | 0.02814 |
| *FCHSD2* | 0.46819 | 0.04708 |
| *FDXR* | 0.93147 | 0.00050 |
| *FGF1* | 0.66769 | 0.00004 |
| *FLII* | 0.46240 | 0.04869 |
| *FNTA* | 0.49563 | 0.04092 |
| *FTO* | 0.33726 | 0.02929 |
| *GAS6* | 0.31437 | 0.00045 |

continued on next page

Supplementary Table S2 – continued from previous page

| Gene | $F_{ST}$ | $p_{F_{ST}}$ |
|---|---|---|
| *GGNBP2* | 0.22363 | 0.03609 |
| *GLCCI1* | 0.57015 | 0.02784 |
| *GRAMD1B* | 0.28781 | 0.01144 |
| *GRM1* | 0.53937 | 0.03379 |
| *GTF3C1* | 0.46246 | 0.04879 |
| *H2AFY2* | 0.22263 | 0.04506 |
| *HSPG2* | 0.34571 | 0.00094 |
| *HTRA1* | 0.22455 | 0.02425 |
| *IK* | 0.79478 | 0.00539 |
| *IP6K1* | 0.35887 | 0.04662 |
| *IPO9* | 0.85115 | 0.00291 |
| *IQSEC1* | 0.38822 | 0.03422 |
| *IQSEC3* | 0.20883 | 0.01999 |
| *KCNK13* | 0.49790 | 0.04120 |
| *KDM2A* | 0.44769 | 0.01671 |
| *KMT2C* | 0.41598 | 0.02534 |
| *KMT5B* | 0.44284 | 0.01807 |
| *LAMP3* | 0.36783 | 0.00367 |
| *LCMT1* | 0.47434 | 0.00262 |
| *LDLR* | 0.22747 | 0.02134 |
| *LLGL2* | 0.47763 | 0.01119 |
| *LPP* | 0.25330 | 0.03910 |
| *LRP11* | 0.64726 | 0.01656 |
| *LSM12* | 0.52516 | 0.03500 |
| *LY96* | 0.42064 | 0.00678 |
| *MAD1L1* | 0.26257 | 0.00400 |
| *MED13L* | 0.29158 | 0.01028 |
| *MED20* | 0.24371 | 0.02413 |
| *MEGF11* | 0.19554 | 0.01331 |
| *MOB3B* | 0.32995 | 0.00299 |
| *MTCL1* | 0.19511 | 0.02164 |
| *MTO1* | 0.42405 | 0.00642 |
| *MYL1* | 0.41734 | 0.00747 |
| *MYO7B* | 0.39156 | 0.01150 |
| *MYO9B** | 0.58992 | 0.02486 |
| *NAA35* | 0.22633 | 0.02278 |
| *NCOR2* | 0.59053 | 0.00209 |
| *NDUFS8* | 0.28450 | 0.01752 |
| *NMUR1* | 0.26947 | 0.03795 |
| *NOL10* | 0.32559 | 0.00224 |
| *NTNG2* | 0.26701 | 0.00112 |
| *NTRK3* | 0.29037 | 0.00293 |
| *NUDT7* | 0.28728 | 0.03905 |
| *NUGGC* | 0.47754 | 0.01010 |
| *NUP93* | 0.23881 | 0.04300 |
| *ODC1* | 0.29926 | 0.00502 |
| *ODF2* | 0.68581 | 0.01385 |
| *PACS1* | 0.33250 | 0.00289 |
| *PANK2* | 0.76970 | 0.00787 |
| *PATJ* | 0.68590 | 0.01376 |
| *PCDH7* | 0.71470 | 0.01159 |
| *PCID2* | 0.97146 | 0.00033 |
| *PDGFRL* | 0.48296 | 0.04418 |
| *PERP* | 0.24191 | 0.03173 |
| *PHC3* | 0.67413 | 0.01424 |

continued on next page

Supplementary Table S2 – continued from previous page

| Gene | $F_{ST}$ | $p_{F_{ST}}$ |
|---|---|---|
| *PLAUR* | 0.22857 | 0.03708 |
| *PLEKHA5* | 0.32369 | 0.01197 |
| *PRDM10* | 0.62931 | 0.01903 |
| *PRKCE* | 0.26969 | 0.00016 |
| *PRRC2A* | 0.33359 | 0.03057 |
| *PTEN* | 0.78622 | 0.00603 |
| *PUM3* | 0.78612 | 0.00612 |
| *PYDC1* | 0.51555 | 0.03711 |
| *RAB11FIP4* | 0.38858 | 0.03391 |
| *RAD51* | 0.36802 | 0.04253 |
| *RBFOX1* | 0.47494 | 0.01168 |
| *RBM33* | 0.26127 | 0.04292 |
| *RFC5* | 0.36285 | 0.04428 |
| *RFX4* | 0.34189 | 0.00013 |
| *RHBDF2* | 0.36213 | 0.00897 |
| *RPS6KA2* | 0.21401 | 0.04906 |
| *RUNDC1* | 0.91873 | 0.00070 |
| *RYR1* | 0.38968 | 0.03413 |
| *SCO1* | 0.64062 | 0.01769 |
| *SDR42E2* | 0.47514 | 0.04549 |
| *SETD3* | 0.40980 | 0.02626 |
| *SLC24A3* | 0.45728 | 0.00364 |
| *SLC45A1* | 0.47461 | 0.04562 |
| *SLC6A13* | 0.35348 | 0.04959 |
| *SPHKAP* | 0.26428 | 0.02997 |
| *SRCAP* | 0.20734 | 0.04156 |
| *SRGN* | 0.51940 | 0.03680 |
| *ST8SIA1* | 0.46287 | 0.04803 |
| *SYT9* | 0.29371 | 0.00615 |
| *TET3* | 0.50453 | 0.03953 |
| *TF* | 0.73293 | 0.01061 |
| *TMEFF2* | 0.31345 | 0.04249 |
| *TMEM178A* | 0.77110 | 0.00755 |
| *TMPRSS9* | 0.54854 | 0.03136 |
| *TNKS* | 0.40762 | 0.00892 |
| *TRBV6-1* | 0.30260 | 0.04972 |
| *TRIM62* | 0.63802 | 0.01795 |
| *TRIM72* | 0.51555 | 0.03821 |
| *TTLL5* | 0.55135 | 0.03075 |
| *UBAC1* | 0.53118 | 0.03537 |
| *UBAC2* | 0.43741 | 0.01960 |
| *UBR3* | 0.37741 | 0.00705 |
| *UMODL1* | 0.33847 | 0.00020 |
| *UQCRC2* | 0.59024 | 0.02441 |
| *URB1* | 0.22569 | 0.01335 |
| *VAC14* | 0.51243 | 0.00138 |
| *WBSCR17* | 0.23422 | 0.03062 |
| *WDFY2* | 0.90447 | 0.00129 |
| *WNT7B* | 0.23625 | 0.03621 |
| *WRAP53* | 0.44955 | 0.01752 |
| *XYLT1* | 0.22870 | 0.00975 |
| *ZBTB3* | 0.41557 | 0.02462 |
| *ZDHHC14* | 0.30653 | 0.00096 |
| *ZFPM2* | 0.26225 | 0.04439 |
| *ZNF3* | 0.33029 | 0.00026 |

continued on next page

Supplementary Table S2 – continued from previous page

| Gene | $F_{ST}$ | $p_{F_{ST}}$ |
|---|---|---|
| *ZNF485* | 0.27073 | 0.00497 |
| *ZNF536* | 0.25102 | 0.03087 |
| *ZNF564* | 0.60466 | 0.02290 |
| *ZSCAN25* | 0.43668 | 0.01962 |
| ENSPANG00000004311 | 0.74849 | 0.00012 |
| ENSPANG00000010457 | 0.61534 | 0.00159 |
| ENSPANG00000017858 | 0.22827 | 0.04616 |
| ENSPANG00000023891 | 0.30260 | 0.04967 |
| ENSPANG00000027984 | 0.65318 | 0.00010 |
| ENSPANG00000028076 | 0.27890 | 0.04559 |
| ENSPANG00000028262 | 0.61153 | 0.00001 |
| ENSPANG00000028279 | 0.42387 | 0.02326 |
| ENSPANG00000028627 | 0.42785 | 0.00637 |

Supplementary Table S3: Genotype frequencies for genes considered to be strong candidates for differentiation (see Table 2). Genome positions and reference alleles are given for the anubis baboon (Panu2.0) genome. The reference allele for the rhesus macaque (Mmul8.0.1) genome was also obtained using a coordinate translation in liftOver (Kent et al., 2002; Rosenbloom et al., 2015). Where the reference alleles differed between genomes, both alleles are listed with the baboon allele listed first. Asterisks (*) indicate that the coordinate could not be converted to Mmul8.0.1, and the reference allele from rheMac2 (an older assembly) is shown instead. Genotype frequencies are shown for each species in the order: homozygous for reference, heterozygous, homozygous for alternate.

| Gene | Position | | Differentiation | | Allelic variants | | Genotype frequencies | | | | | |
|---|---|---|---|---|---|---|---|---|---|---|---|---|
| | (Panu2.0) | | $F_{ST}$ | $p_{F_{ST}}$ | Ref. | Alt. | Kinda | | | chacma | | |
| *ATXN2* | chr11 | 110620892 | 0.84882 | 0.00003 | G | A | 54 | 13 | 0 | 0 | 0 | 15 |
| | chr11 | 110620934 | 0.84882 | | A | G | 0 | 13 | 54 | 15 | 0 | 0 |
| *EFHD2* | chr1 | 17875364 | 0.61707 | < 0.00001 | C | G | 64 | 2 | 0 | 4 | 3 | 2 |
| | chr1 | 17875397 | 0.71940 | | T | C | 66 | 0 | 0 | 4 | 3 | 2 |
| | chr1 | 17875411 | 0.71940 | | T | G | 66 | 0 | 0 | 4 | 3 | 2 |
| | chr1 | 17875416 | -0.00265 | | G/A | A | 9 | 57 | 0 | 3 | 3 | 3 |
| | chr1 | 17875436 | 0.71940 | | C/T | T | 66 | 0 | 0 | 4 | 3 | 2 |
| | chr1 | 17875441 | 0.71940 | | C | T | 66 | 0 | 0 | 4 | 3 | 2 |
| *FGF1* | chr6 | 136137340 | 0.23022 | 0.00004 | T/T* | C | 12 | 25 | 26 | 0 | 1 | 15 |
| | chr6 | 136137374 | 0.87739 | | T/T* | C | 62 | 1 | 0 | 2 | 3 | 11 |
| | chr6 | 136137439 | 0.89546 | | T/T* | G | 62 | 1 | 0 | 1 | 4 | 11 |
| *PRKCE* | chr13 | 44855469 | 0.51137 | 0.00016 | A/G | G | 6 | 25 | 36 | 10 | 0 | 1 |
| | chr13 | 44855470 | 0.51137 | | C | G | 6 | 25 | 36 | 10 | 0 | 1 |
| | chr13 | 44855471 | 0.51137 | | A | G | 6 | 25 | 36 | 10 | 0 | 1 |
| | chr13 | 44855507 | 0.51137 | | G/C | C | 6 | 25 | 36 | 10 | 0 | 1 |
| | chr13 | 44898572 | 0.71643 | | A | C | 66 | 0 | 0 | 5 | 4 | 4 |
| | chr13 | 44898586 | -0.01417 | | C | T | 65 | 2 | 0 | 13 | 0 | 0 |
| | chr13 | 44898587 | 0.52273 | | G | A | 61 | 6 | 0 | 5 | 4 | 4 |
| | chr13 | 44898613 | 0.51940 | | G | A | 67 | 0 | 0 | 8 | 3 | 2 |
| | chr13 | 44898617 | 0.51940 | | G | A | 67 | 0 | 0 | 8 | 3 | 2 |
| | chr13 | 44898696 | -0.00921 | | T | A | 63 | 3 | 1 | 13 | 0 | 0 |
| | chr13 | 45024148 | -0.01925 | | A | C | 57 | 7 | 1 | 12 | 1 | 0 |
| | chr13 | 45024166 | -0.01605 | | G | A | 62 | 2 | 1 | 13 | 0 | 0 |
| | chr13 | 45024206 | 0.71375 | | A | C | 65 | 0 | 0 | 5 | 4 | 4 |
| | chr13 | 45024265 | -0.01925 | | C/T | T | 57 | 7 | 1 | 12 | 1 | 0 |
| | chr13 | 45053068 | 0.17177 | | C | T | 68 | 0 | 0 | 12 | 0 | 1 |
| | chr13 | 45053077 | 0.08873 | | C | T | 67 | 1 | 0 | 12 | 0 | 1 |
| | chr13 | 45053155 | 0.17177 | | G | A | 68 | 0 | 0 | 12 | 0 | 1 |
| | chr13 | 45053159 | 0.17177 | | G | C | 68 | 0 | 0 | 12 | 0 | 1 |
| | chr13 | 45053165 | 0.11253 | | T | G | 44 | 20 | 4 | 13 | 0 | 0 |
| | chr13 | 45150212 | 0.00257 | | C | T | 55 | 4 | 0 | 16 | 0 | 0 |
| | chr13 | 45150236 | -0.01482 | | C | T | 58 | 1 | 0 | 16 | 0 | 0 |

continued on next page

Supplementary Table S3 – continued from previous page

| Gene | Position (Panu2.0) | | Differentiation $F_{ST}$ | $p_{F_{ST}}$ | Allelic variants Ref. | Alt. | Genotype frequencies Kinda | | | chacma | | |
|---|---|---|---|---|---|---|---|---|---|---|---|---|
| *RFX4* | chr11 | 105851669 | 0.72300 | 0.00013 | C | T | 2 | 13 | 51 | 11 | 3 | 0 |
| | chr11 | 105851730 | -0.01749 | | A | G | 65 | 1 | 0 | 14 | 0 | 0 |
| | chr11 | 105870425 | 0.68939 | | C | T | 65 | 0 | 0 | 5 | 4 | 3 |
| | chr11 | 105870454 | -0.01573 | | C | T | 63 | 2 | 0 | 12 | 0 | 0 |
| | chr11 | 105870471 | 0.44224 | | C | A | 66 | 0 | 0 | 9 | 1 | 2 |
| | chr11 | 105870476 | 0.09165 | | T | C | 44 | 21 | 1 | 12 | 0 | 0 |
| | chr11 | 105870506 | 0.44224 | | T | G | 66 | 0 | 0 | 9 | 1 | 2 |
| | chr11 | 105870520 | 0.44224 | | C | T | 66 | 0 | 0 | 9 | 1 | 2 |
| | chr11 | 105870526 | 0.44224 | | A | G | 66 | 0 | 0 | 9 | 1 | 2 |
| | chr11 | 105870586 | 0.44224 | | C | T | 66 | 0 | 0 | 9 | 1 | 2 |
| | chr11 | 105870588 | 0.07881 | | C | G | 48 | 14 | 4 | 12 | 0 | 0 |
| *UMODL1* | chr3 | 4512016 | 0.40211 | 0.00020 | A/T | G | 68 | 1 | 0 | 10 | 0 | 3 |
| | chr3 | 4512024 | 0.46734 | | T/A | G | 69 | 0 | 0 | 10 | 0 | 3 |
| | chr3 | 4512027 | 0.46734 | | A/T | G | 69 | 0 | 0 | 10 | 0 | 3 |
| | chr3 | 4512050 | 0.46734 | | G/C | C | 69 | 0 | 0 | 10 | 0 | 3 |
| | chr3 | 4512065 | 0.06414 | | C/A | T | 52 | 15 | 2 | 13 | 0 | 0 |
| | chr3 | 4512066 | 0.01904 | | G/C | A | 68 | 1 | 0 | 13 | 0 | 0 |
| | chr3 | 4512075 | 0.00457 | | C/A | T | 63 | 6 | 0 | 13 | 0 | 0 |
| | chr3 | 4512076 | 0.46734 | | G/C | A | 69 | 0 | 0 | 10 | 0 | 3 |
| | chr3 | 4512080 | 0.46734 | | C/G | T | 69 | 0 | 0 | 10 | 0 | 3 |
| | chr3 | 4512092 | 0.46734 | | A/T | G | 69 | 0 | 0 | 10 | 0 | 3 |
| | chr3 | 4512098 | 0.46734 | | T/A | C | 69 | 0 | 0 | 10 | 0 | 3 |
| ENSPANG-00000004311 | chr10 | 86441991 | 0.74849 | 0.00012 | G | A | 50 | 16 | 2 | 0 | 1 | 13 |
| | chr10 | 86441993 | 0.74849 | | T | A | 50 | 16 | 2 | 0 | 1 | 13 |
| ENSPANG-00000027984 | chr13 | 5031394 | 0.75948 | 0.00010 | C | G | 58 | 7 | 2 | 2 | 1 | 12 |
| | chr13 | 5031410 | 0.44058 | | G | A | 67 | 0 | 0 | 10 | 3 | 2 |
| | chr13 | 5031434 | 0.75948 | | T | A | 58 | 7 | 2 | 2 | 1 | 12 |
| ENSPANG-00000028262 | chr1 | 185450553 | 0.19649 | 0.00001 | A | G | 65 | 1 | 0 | 13 | 0 | 2 |
| | chr1 | 185450577 | 0.86013 | | G | A | 66 | 0 | 0 | 3 | 3 | 9 |
| | chr1 | 185450607 | 0.57834 | | C | T | 52 | 12 | 2 | 3 | 2 | 9 |
| | chr1 | 185450667 | 0.81117 | | T | C | 66 | 0 | 0 | 4 | 3 | 7 |

Supplementary Table S4: Gene Ontology (GO) terms (biological process ontology) with significantly enriched differentiation. Terms that are enriched exhibit an overall shift in $p_{F_{ST}}$. Only terms with $p$ < 0.05 are shown here.

| GO term | | Enrichment ($p$-value) |
|---|---|---|
| Accession | Name | |
| GO:0045598 | regulation of fat cell differentiation | 0.00009 |
| GO:0015850 | organic hydroxy compound transport | 0.00062 |
| GO:0048639 | positive regulation of developmental growth | 0.00099 |
| GO:0045839 | negative regulation of mitotic nuclear division | 0.00130 |
| GO:0061448 | connective tissue development | 0.00222 |
| GO:0050772 | positive regulation of axonogenesis | 0.00230 |
| GO:0001936 | regulation of endothelial cell proliferation | 0.00357 |
| GO:0031667 | response to nutrient levels | 0.00383 |
| GO:0050680 | negative regulation of epithelial cell proliferation | 0.00431 |
| GO:0048589 | developmental growth | 0.00826 |
| GO:0002062 | chondrocyte differentiation | 0.00864 |
| GO:0006816 | calcium ion transport | 0.00927 |
| GO:0000122 | negative regulation of transcription by RNA polymerase II | 0.01012 |
| GO:0034968 | histone lysine methylation | 0.01084 |
| GO:0051051 | negative regulation of transport | 0.01102 |
| GO:0097035 | regulation of membrane lipid distribution | 0.01320 |
| GO:0046677 | response to antibiotic | 0.01501 |
| GO:0035264 | multicellular organism growth | 0.01546 |
| GO:0051345 | positive regulation of hydrolase activity | 0.01546 |
| GO:0051304 | chromosome separation | 0.01547 |
| GO:0051896 | regulation of protein kinase B signaling | 0.01583 |
| GO:0051209 | release of sequestered calcium ion into cytosol | 0.01590 |
| GO:0007600 | sensory perception | 0.01612 |
| GO:1901565 | organonitrogen compound catabolic process | 0.01622 |
| GO:0043062 | extracellular structure organization | 0.01628 |
| GO:0010717 | regulation of epithelial to mesenchymal transition | 0.01635 |
| GO:0061136 | regulation of proteasomal protein catabolic process | 0.01748 |
| GO:0090630 | activation of GTPase activity | 0.01779 |
| GO:0006413 | translational initiation | 0.01833 |
| GO:0090316 | positive regulation of intracellular protein transport | 0.01949 |
| GO:0010001 | glial cell differentiation | 0.02024 |
| GO:0045444 | fat cell differentiation | 0.02044 |
| GO:0001659 | temperature homeostasis | 0.02143 |
| GO:0051240 | positive regulation of multicellular organismal process | 0.02185 |
| GO:0072006 | nephron development | 0.02412 |
| GO:0051983 | regulation of chromosome segregation | 0.02472 |
| GO:0007417 | central nervous system development | 0.02552 |
| GO:0062012 | regulation of small molecule metabolic process | 0.02644 |
| GO:0009636 | response to toxic substance | 0.02648 |
| GO:0035148 | tube formation | 0.02723 |
| GO:0006259 | DNA metabolic process | 0.02780 |
| GO:0051054 | positive regulation of DNA metabolic process | 0.02968 |
| GO:1901991 | negative regulation of mitotic cell cycle phase transition | 0.03032 |
| GO:0000070 | mitotic sister chromatid segregation | 0.03134 |
| GO:0008361 | regulation of cell size | 0.03193 |
| GO:0030307 | positive regulation of cell growth | 0.03522 |
| GO:0048259 | regulation of receptor-mediated endocytosis | 0.03574 |
| GO:0120161 | regulation of cold-induced thermogenesis | 0.03639 |
| GO:0048675 | axon extension | 0.03649 |
| GO:0015914 | phospholipid transport | 0.03711 |

continued on next page

Supplementary Table S4 – continued from previous page

| GO term | | Enrichment ($p$-value) |
|---|---|---|
| Accession | Name | |
| GO:0048878 | chemical homeostasis | 0.03773 |
| GO:0010906 | regulation of glucose metabolic process | 0.03791 |
| GO:0050801 | ion homeostasis | 0.04018 |
| GO:0051050 | positive regulation of transport | 0.04124 |
| GO:0031328 | positive regulation of cellular biosynthetic process | 0.04193 |
| GO:0080090 | regulation of primary metabolic process | 0.04213 |
| GO:0000724 | double-strand break repair via homologous recombination | 0.04321 |
| GO:0001568 | blood vessel development | 0.04374 |
| GO:1901361 | organic cyclic compound catabolic process | 0.04429 |
| GO:0050921 | positive regulation of chemotaxis | 0.04542 |
| GO:0007411 | axon guidance | 0.04563 |
| GO:0050796 | regulation of insulin secretion | 0.04675 |
| GO:0097306 | cellular response to alcohol | 0.04691 |
| GO:0006302 | double-strand break repair | 0.04760 |
| GO:0032956 | regulation of actin cytoskeleton organization | 0.04812 |
| GO:0006939 | smooth muscle contraction | 0.04881 |

Supplementary Table S5: $F_{ST}$ and $p$ values of genes in the CCKR signaling pathway. Genome positions are given for the anubis baboon (Panu2.0) genome.

| Gene | | Position | $F_{ST}$ | $p_{F_{ST}}$ | BH-adjusted $p_{F_{ST}}$ |
|---|---|---|---|---|---|
| *GNB1* | chr1 | 4724905-4839528 | 0.12757 | 0.34870 | > 0.99999 |
| *ODC1* | chr1 | 33711839-33721404 | 0.29926 | 0.00502 | 0.27773 |
| *FOXO3* | chr4 | 153548329-153654266 | 0.02159 | 0.87560 | > 0.99999 |
| *PIK3R1* | chr6 | 63980911-64062226 | 0.05827 | 0.76380 | > 0.99999 |
| *LYN* | chr8 | 51971047-52057050 | 0.12235 | 0.46480 | > 0.99999 |
| *PTEN* | chr9 | 80021075-80056543 | 0.78622 | 0.00603 | 0.30886 |
| *PRKCE* | chr13 | 44756709-45175428 | 0.26969 | 0.00016 | 0.03934 |
| *TPCN2* | chr14 | 5295886-5334864 | 0.22514 | 0.06081 | 0.65791 |
| *GUCY2D* | chr16 | 7707896-7725235 | 0.07230 | 0.61890 | > 0.99999 |
| *AKAP1* | chr16 | 38017354-38051906 | 0.02482 | 0.70890 | > 0.99999 |
| *STAT3* | chr16 | 48978970-49012645 | 0.11931 | 0.36770 | > 0.99999 |
| *FOXO1* | chr17 | 19194223-19301111 | 0.17917 | 0.22320 | 0.95342 |
| *MAP2K2* | chr19 | 3900243-3938641 | 0.24289 | 0.08564 | 0.76623 |
| *RYR1* | chr19 | 32699970-32857343 | 0.38968 | 0.03413 | 0.58253 |

Supplementary Table S6: Genotype frequencies for all components and genes in the JAK/STAT signaling pathway with identified variants in our dataset. Genome positions and reference alleles are given for the anubis baboon (Panu2.0) genome. The reference allele for the rhesus macaque (Mmul8.0.1) genome was also obtained using a coordinate translation in liftOver (Kent et al., 2002; Rosenbloom et al., 2015). Where the reference alleles differed between genomes, both alleles are listed with the baboon allele listed first. Genotype frequencies are shown for each species in the order: homozygous for reference, heterozygous, homozygous for alternate.

| Component | Gene | Position (Panu2.0) | | Differentiation $F_{ST}$ | $p_{F_{ST}}$ | Allelic variants Ref. | Alt. | Genotype frequencies Kinda | | | chacma | | |
|---|---|---|---|---|---|---|---|---|---|---|---|---|---|
| JAK | *JAK1* | chr1 | 66951070 | 0.07908 | 0.14740 | A | C | 65 | 0 | 0 | 18 | 0 | 1 |
| | | chr1 | 66951074 | 0.07908 | | C | A | 65 | 0 | 0 | 18 | 0 | 1 |
| | | chr1 | 66951078 | 0.07908 | | C | G | 65 | 0 | 0 | 18 | 0 | 1 |
| | | chr1 | 66951082 | 0.59910 | | T/C | C | 0 | 3 | 62 | 6 | 7 | 6 |
| PIAS | *PIAS1* | chr7 | 42548519 | 0.39690 | 0.06619 | A | T | 54 | 12 | 2 | 4 | 4 | 5 |
| | *PIAS4* | chr19 | 3844945 | 0.21996 | 0.17450 | C | T | 67 | 0 | 0 | 12 | 3 | 0 |
| STAT | *STAT3* | chr16 | 48997534 | 0.11931 | 0.36770 | C | T | 59 | 0 | 0 | 14 | 2 | 0 |
